# Neurodevelopmental defects in a mouse model of O-GlcNAc transferase intellectual disability

**DOI:** 10.1101/2023.08.23.554427

**Authors:** Florence Authier, Nina Ondruskova, Andrew T. Ferenbach, Alison McNeilly, Daan M. F. van Aalten

**Affiliations:** Department of Molecular Biology and Genetics, Aarhus University, Aarhus, Denmark; Department of Paediatrics and Inherited Metabolic Disorders, First Faculty of Medicine, Charles University and General University Hospital in Prague, Prague, Czech Republic; Division of Systems Medicine, School of Medicine, University of Dundee, Dundee, UK; Division of Cell and Developmental Biology, School of Life Sciences, University of Dundee, Dundee, UK

## Abstract

O-GlcNAcylation is a protein modification that is critical for vertebrate development, catalysed by O-GlcNAc transferase (OGT) and reversed by O-GlcNAcase (OGA). Missense mutations in *OGT* have recently been shown to segregate with a syndromic form of intellectual disability, OGT-linked Congenital Disorder of Glycosylation (OGT-CDG). Although OGT-CDG suggests a critical role of O-GlcNAcylation in neurodevelopment and/or cognitive function, the underlying pathophysiologic mechanisms remain unknown. Here we report three mouse lines that carry three different catalytically impaired OGT-CDG variants. These mice show altered O-GlcNAc homeostasis with decreased global O-GlcNAcylation and OGT/OGA levels in the brain. Phenotypic characterization of the mice revealed microcephaly and cognitive deficits including hyperactivity, anxiety and altered spatial working memory. These mouse models will serve as an important tool to study genotype-phenotype correlation in OGT-CDG *in vivo* and for the development of possible treatment avenues for this disorder.

**Significant statement:** Mutations in O-GlcNAc transferase (OGT), the sole enzyme that installs O-GlcNAc sugar on proteins, lead to intellectual disability through unknown mechanisms. We have generated mouse models carrying OGT mutations that show reduction in brain size, hyperactivity and defects in memory. These mouse models will serve as a valuable tool to further investigate disease mechanism and propose future treatment avenues.

## Introduction

Intellectual disability (ID) represents a heterogeneous group of neurodevelopmental disorders which are predicted to affect up to 1-3% of the worldwide population (1). ID appears during childhood and is characterized by impaired cognitive function (with IQ below 70) and adaptive behaviour (2). Frequent comorbidities are autism spectrum disorders (ASD), attention deficit and hyperactivity disorder (ADHD) and epilepsy (3).

O-GlcNAcylation is a dynamic and highly conserved posttranslational modification of serine and threonine residues on proteins that regulate several cellular processes including transcription (4,5), signalling (6) and metabolism (7). Only two enzymes regulate O-GlcNAcylation: O-GlcNAc transferase (OGT) is responsible for the addition of a single *N*-acetyl glucosamine (GlcNAc) moiety on substrates (8) and O-GlcNAcase (OGA) for its removal (9). O-GlcNAcylation has been shown to be essential in mammals. *Ogt* deletion leads to impaired embryogenesis and early development (10,11) while *Oga* is critical for perinatal survival (12–14). Deregulation of O-GlcNAcylation homeostasis has also been associated with pathological conditions such as diabetes, neurodegeneration and cancers (15–17).

O-GlcNAcylation and the two cycling enzymes are abundant in the mammalian brain (9,18,19), in particular at pre- and postsynaptic compartments where many proteins important for neuronal structure and synaptic function have been found to be O-GlcNAcylated (20–23). Previous work has demonstrated that modulation of O-GlcNAcylation affects neuronal processes important for brain function, including synaptic maturation, synaptic function and axon morphology and learning and memory (22,24–28). While collectively these suggest an important role for neuronal O-GlcNAcylation in cognition, our understanding of how O-GlcNAcylation and the cycling enzymes regulate brain function is still limited.

Recently, missense mutations in *OGT* have been identified in patients affected with intellectual disability, giving rise to a syndrome named OGT-linked Congenital Disorder of Glycosylation (OGT-CDG) (29). As *OGT* is located on the X chromosome, almost all patients are male although a *de novo* missense variant has also been reported in a monozygotic female twin (30). OGT-CDG is a clinically heterogenous disorder in which all patients present with intellectual disability, developmental delay and very restricted language skills. OGT-CDG patients also commonly present with dysmorphic features including craniofacial characteristics with broad and high forehead, hypertelorism, broad nasal root, full or long philtrum, and clinodactyly. Brain and eye abnormalities are also observed in most patients (29).

OGT is composed of a catalytic domain and a N-terminal tetratricopeptide repeat (TPR) domain consisting of 13.5 TPRs responsible for substrate binding and protein:protein interactions (31). In addition to installing O-GlcNAc on to proteins, OGT is involved in the proteolytic processing and activation of the host cell factor 1 (HCF1) (32), a known ID gene (33) and possesses non-catalytic functions implicated in cellular proliferation (34). To date, 17 OGT-CDG variants have been reported in *OGT* with mutations in the catalytic and TPR domains giving rise to similar clinical features. This suggest that there are common mechanisms affected in these OGT-CDG variants, which remain unknown as no vertebrate models are currently available to dissect these. Investigating the function of OGT in neurodevelopment and the brain has been hampered by mouse lethality caused by global loss of *Ogt* (10,11). Although several brain cell specific *Ogt* knock out (KO) mouse models also lead to postnatal lethality, these have highlighted the role of OGT in neurodevelopment, neuronal survival and structure (11,28,35–37). Modelling OGT-CDG *in vivo* will help to more precisely understand how OGT regulates processes that are essential for neurodevelopment and brain function.

Here, we report the use of a CrisprCas/9 genome editing approach to generate three mouse models carrying three different catalytically impaired OGT-CDG variants: C921Y, N648Y and N567K. Unlike previous OGT KO models, OGT-CDG mice are viable allowing the phenotypic characterization of the animals. Loss of OGT catalytic activity leads to impaired O-GlcNAcylation homeostasis in the brain and changes in size and mass of OGT-CDG mice. In addition, OGT^C921Y^ mice show microcephaly and cognition deficits including hyperactivity, anxiety and memory defects, recapitulating clinical OGT-CDG symptoms.

## Results

### OGT-CDG mutant mice are viable

To dissect the effects of OGT-CDG mutations *in vivo*, we used a CRISPR/Cas9 genome editing approach in mice to introduce the missense variants C921Y (38), N648Y (39) and N567K (30), all reported previously and located in the OGT catalytic domain (**Fig. 1A**). Briefly, 0.5 dpc zygotes were injected with editing reagents and transferred into pseudo-pregnant female mice. DNA from offspring was genotyped and sequenced to confirm the presence of the OGT-CDG mutations for each of the three mouse lines (**Fig. 1B, C and D**). As *Ogt* is essential for embryogenesis (10,11), we first assessed the viability and mendelian inheritance distribution of offspring generated from heterozygous OGT^CDG/+^ females crossed to OGT^WT^ males on the same C57BL/6J genetic background. This generated 48.8%, 58.1% and 46.5% of male pups for the OGT^C921Y^, OGT^N648Y^, OGT^N567K^ lines, respectively and 51.2%, 41.9% and 53.5% of female pups for the OGT^C921Y^, OGT^N648Y^ and OGT^N567K^ lines respectively. These suggest no gender-related lethality in these OGT-CDG mouse lines. DNA from offspring was then genotyped and sequenced, revealing that both heterozygous female and hemizygous male offspring could be obtained for all three OGT-CDG lines. We then analysed mendelian inheritance of wild type OGT^WT^ (WT) (expected 50%), heterozygous OGT^CDG/+^ (expected 25%) and hemizygous OGT^CDG^ (expected 25%) animals arising from all the lines. For the OGT^C921Y^ line, this generated 27.3% of WT female, 23.8% of OGT^C921Y/+^ female, 23.8% of WT male and 25% OGT^C921Y^ male at the expected mendelian ratio (Chi-square = 0.29, n = 84). For the OGT^N648Y^ line, this generated 25.8% of OGT^N648Y/+^ female, 29% WT male and 29% OGT^N648Y^ male at expected ratio (Chi-square = 1.39, n = 31) while a slight reduced percentage of WT female (16.1%) pups were born. Intriguingly, although no significant differences were observed in litter size in the OGT^N567K^ line compared to OGT^C921Y^ and OGT^N648Y^ lines, we observed a significant deviation from the expected mendelian ratio (Chi-square = 12.35, n = 43, \*\**p* = 0.006) where 44.2% of WT female, 30.2% of WT male and only 9.3% of OGT^N567K/+^ female and 16.3% of OGT^N567K^ male were genotyped after weaning (**Fig. 1E**). These suggest that the N567K mutation may affect the genotype distribution among offspring – it should be noted that this *de novo* variant has only been found in female twins with skewed X-inactivation (30). We also investigated the fertility of male hemizygotes carrying OGT-CDG mutations as some of the patients affected with OGT-CDG exhibit genital abnormalities (29). We have successfully obtained WT and heterozygous OGT-CDG animals from WT female and hemizygous OGT-CDG male breeding pairs for all three lines indicating that fertility is not affected in OGT-CDG mice. As the breeding of the OGT^C921Y^ line generated a sufficient number of animals with expected Mendelian ratio, we performed further detailed phenotypic characterizations on this line. Data for both OGT^N648Y^ and OGT^N567K^ lines, when available, are included in the supplemental section. Taken together, these results show that contrary to *Ogt* KO models, animals carrying OGT-CDG mutations in the catalytic domain of OGT are viable.

**Figure 1:**
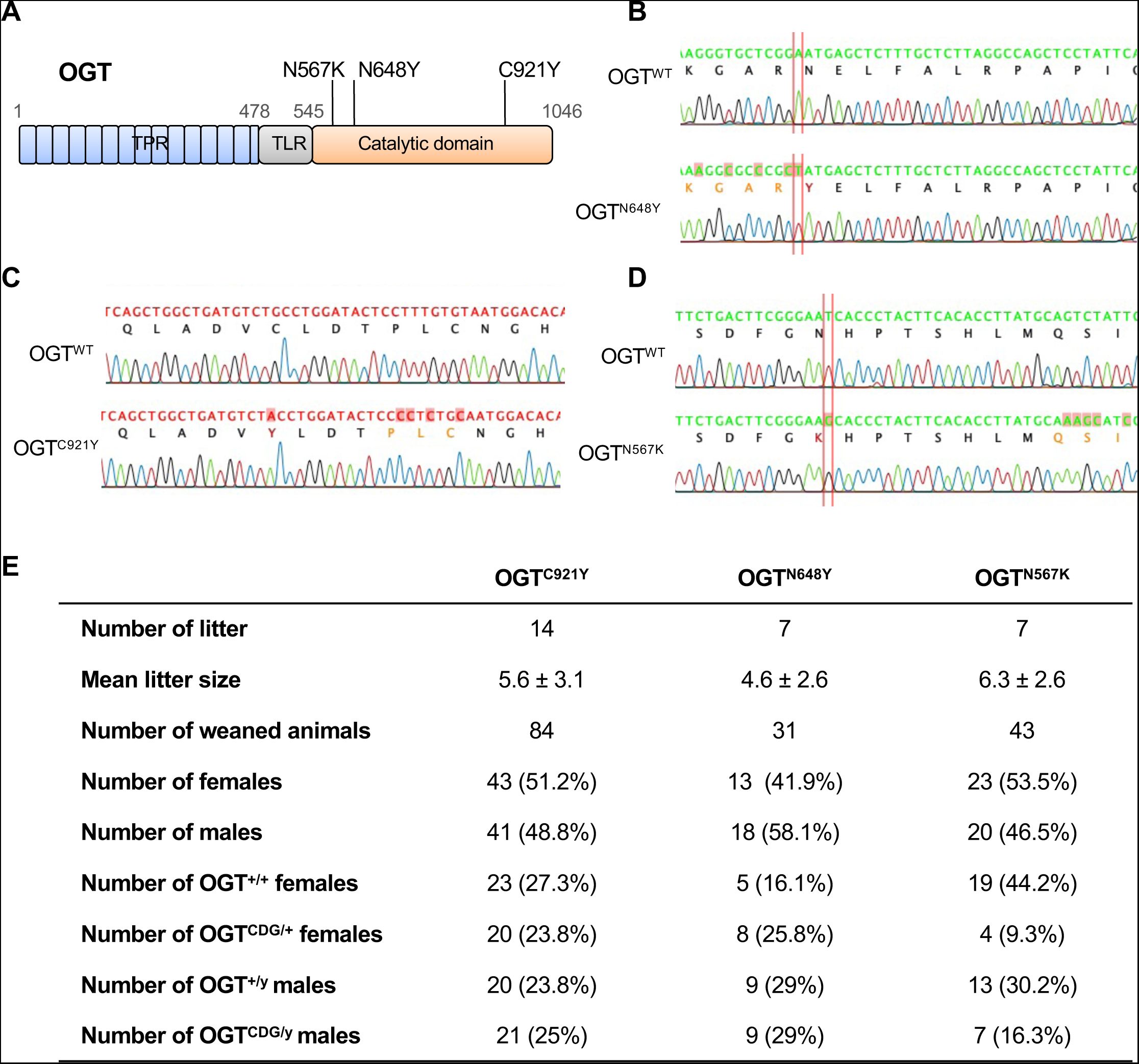
Genome editing to introduce OGT-CDG mutations leads to viable mice. **(a)** Schematic of the OGT protein with three OGT-CDG variants. TPR (blue), TLR (grey) and catalytic (orange) domains are represented. **(b)** Sequencing of genomic DNA of WT and OGT^C921Y^ confirms the presence of the C921Y point mutation (in red) in the transgenic animals. **(c)** Sequencing of genomic DNA of WT and OGT^N648Y^ confirms the presence of the N648Y point mutation (in red) in the transgenic animals. **(d)** Sequencing of genomic DNA of WT and OGT^N567K^ confirms the presence of the N567K point mutation (in red) in the transgenic animals. **(e)** Table showing numbers and percentages of female OGT^+/+^, male OGT^+/y^, female OGT^CDG/+^ and male OGT^CDG/y^ animals generated from female OGT^CDG/+^ x male OGT^+/y^ breeding pairs.

### OGT-CDG mutations are linked to changes in brain O-GlcNAc homeostasis

The OGT^C921Y^ variant affects the catalytic activity of OGT, leading to disruption of O-GlcNAcylation homeostasis in stem cells (38). Therefore, we first investigated the levels of OGT, OGA and O-GlcNAc levels in whole brains from OGT^C921Y^ animals (**Fig. 2**). Global protein O-GlcNAc levels were significantly decreased in OGT^C921Y^ hemizygous males compared to their WT littermates (**Fig. 2A, B**). In addition, both OGA (**Fig. 2A, C**) and OGT (**Fig. 2A, D**) protein levels were also significantly reduced in OGT^C921Y^ animals compared to WT. To investigate whether reduction in OGT and OGA protein levels were due to reduced transcription, we performed qPCR analysis on whole brain tissue from WT and OGT^C921Y^ mice. We observed a reduction in *Oga* mRNA levels in OGT^C921Y^ animals (**Fig. 2E**), accompanied by an increase in *Ogt* mRNA levels (**Fig. 2F**). As modulation of O-GlcNAc levels has been shown to affect splicing of detained introns of *Ogt* and therefore mRNA abundance (40), we performed qPCR analysis using primers specific for detained introns (DI) and decoy exon (PE) *Ogt* transcripts. We have observed a decrease in both detained introns and decoy exon levels transcripts (**Fig. S2**), suggesting that the reduction in OGT protein levels is not caused by alternative splicing but occurs at the protein level. Similar O-GlcNAcylation homeostasis disruption and changes in *Oga* and *Ogt* mRNA levels were observed in the brain of hemizygous OGT^N567K^ and OGT^N648Y^ animals compared to their WT counterparts (**Fig. S1, S2**). Taken together, these results show that OGT-CDG mutations are linked to changes in brain O-GlcNAc homeostasis.

**Figure 2:**
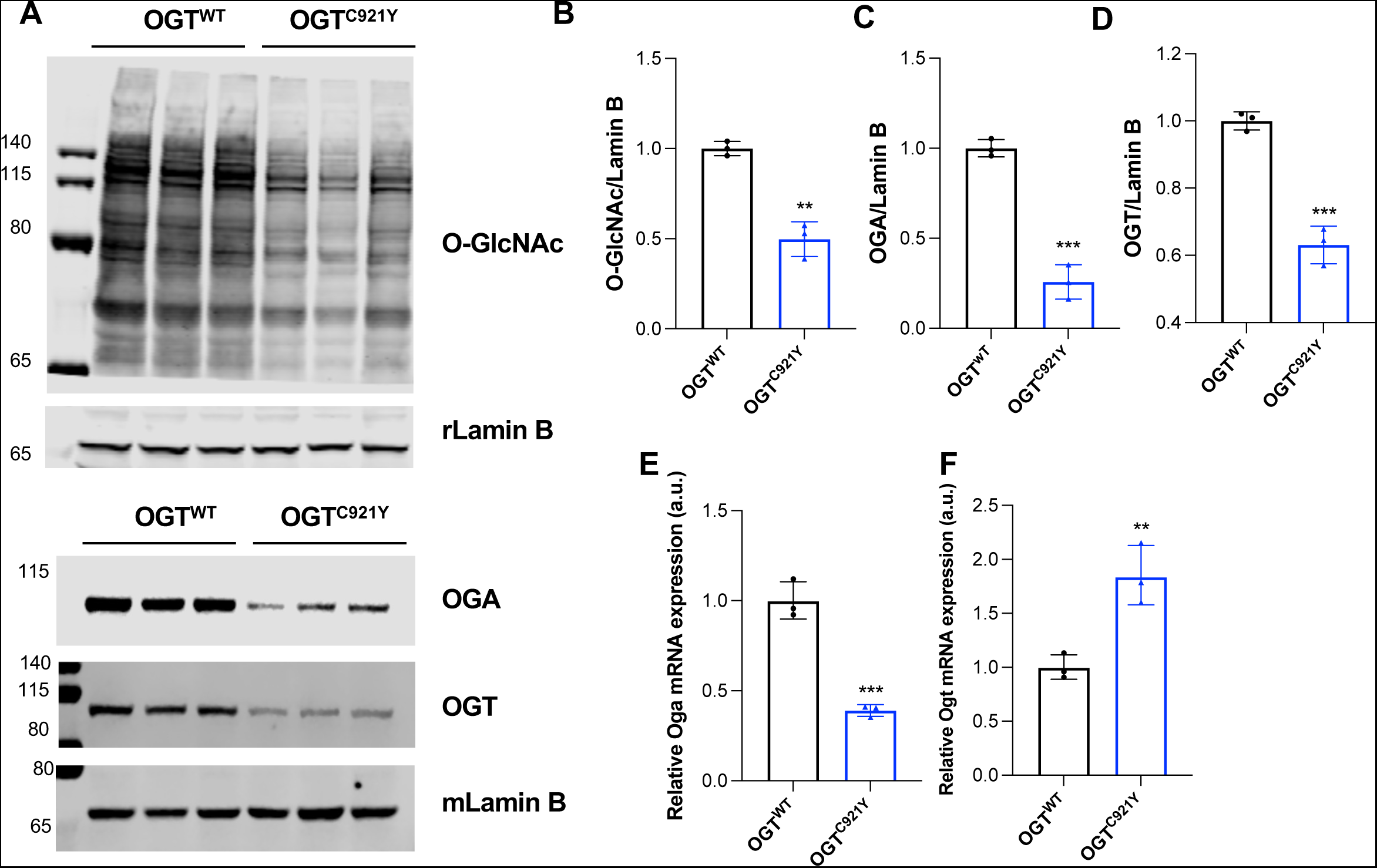
OGT^C921Y^ mutation causes changes in brain O-GlcNAc homeostasis. Data are represented as mean ± SD, n = 3 for all genotypes. Significance is shown as \**p* < 0.05, \*\**p* < 0.01 and \*\*\**p* < 0.001. **(a)** Western blot of O-GlcNAc, OGA and OGT levels in adult brain of OGT^WT^ and OGT^C921Y^ mice. Rabbit and mouse Lamin B antibodies were used as loading control. **(b)** Quantification of O-GlcNAcylated proteins from the Western blot in panel a). **(c)** Quantification of OGA proteins levels from the Western blot in panel a). **(d)** Quantification of OGT proteins levels from the Western blot in panel a). **(e)** Quantification of *Oga* mRNA levels in whole mouse brain by RT-PCR. **(f)** Quantification of *Ogt* mRNA levels in whole mouse brain by RT-PCR.

### OGT-CDG mutations affect mouse size, weight and fat mass

OGT-CDG patients commonly present with short stature and low birth weight and both features were also observed in individuals carrying the OGT^C921Y^ variant (29,38). We therefore evaluated morphometric parameters in OGT^C921Y^ animals. We observed a significant decrease in both body weight (**Fig. 3A**) and nose-to-tail length (**Fig. 3B**) in OGT^C921Y^ mice compared to their WT littermates. Similarly, significantly reduced body weight (**Fig. S3A, B**) and nose-to-tail length (**Fig. S2D, E**) were observed in OGT^N648Y^ and OGT^N567K^ mouse lines. These quantitative changes were accompanied with a slim appearance of mutant animals in all three lines. We next performed EchoMRI imaging to evaluate body composition in the OGT^C921Y^ mice. We observed a significant reduction in body fat mass in OGT^C921Y^ mice compared to WT animals (**Fig. 3C**). These mice also showed an increase in lean body mass (**Fig. 3D**). This suggests that the reduction in body weight observed in OGT-CDG mice is due to reduced adipose tissue content as well as short stature. In addition, OGT^C921Y^ mice display lower levels of glycemia (**Fig. 3E**) compared to WT, suggesting that these mice may possess metabolic phenotypes. Interestingly, lower glycemia was combined with a significant increase in pancreas weight in OGT^C921Y^ animals compared to WT (**Table 1**). Similar change in pancreas weight were observed in both OGT^N648Y^ and OGT^N567K^ animals (**Table 1**). Taken together, these results suggest that OGT-CDG mutations affect mouse size, weight and fat mass.

**Figure 3:**
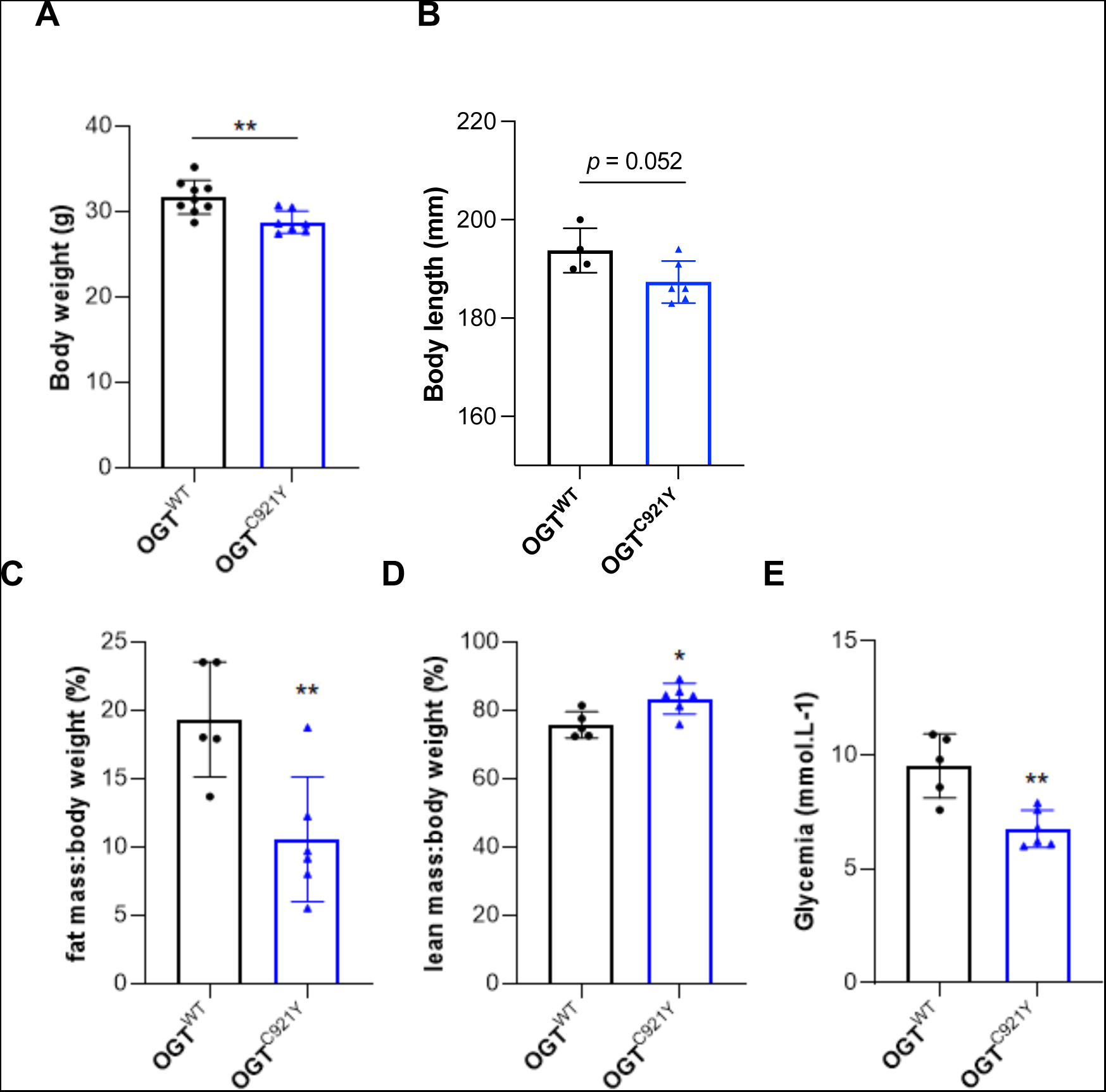
OGT^C921Y^ mutation leads to changes in mass and size. Data are represented as mean ± SD. Significance is shown as \**p* < 0.05, \*\**p* < 0.01 and \*\*\**p* < 0.001. **(a)** Measurement of body weight of OGT^WT^ (n = 9) and OGT^C921Y^ (n = 7) mice. **(b)** Measurement of body length (nose to tail length) of OGT^WT^ (n = 4) and OGT^C921Y^ (n = 6) mice. **(c)** Percentage of fat mass:body weight ratio of OGT^WT^ (n = 5) and OGT^C921Y^ (n = 6) mice. **(d)** Percentage of lean mass:body weight ratio of OGT^WT^ (n = 5) and OGT^C921Y^ (n = 6) mice. **(e)** Basal glycemia levels of OGT^WT^ (n = 5) and OGT^C921Y^ (n = 6) mice.

**Table 1:** Tissue weight analysis in OGT-CDG mice.

|  |  | OGT<sup>WT</sup> | OGT<sup>C921Y</sup> | OGT<sup>WT</sup> | OGT<sup>N648Y</sup> | OGT<sup>WT</sup> | OGT<sup>N567K</sup> |
| --- | --- | --- | --- | --- | --- | --- | --- |
| **Thymus** | Abs (g) | 0.08 ± 0.02 | 0.06 ± 0.01* | 0.1 ± 0.03 | 0.08 ± 0.01 | 0.06 ± 0.03 | 0.05 ± 0.02 |
|  | % of BW | 0.25 ± 0.05 | 0.21 ± 0.02 | 0.28 ± 0.09 | 0.27 ± 0.03 | 0.20 ± 0.07 | 0.19 ± 0.06 |
| **Heart** | Abs (g) | 0.25 ± 0.05 | 0.23 ± 0.04 | 0.22 ± 0.01 | 0.24 ± 0.03 | 0.26 ± 0.06 | 0.23 ± 0.02 |
|  | % of BW | 0.77 ± 0.15 | 0.80 ± 0.14 | 0.66 ± 0.04 | 0.82 ± 0.04** | 0.83 ± 0.16 | 0.84 ± 0.08 |
| **Lungs** | Abs (g) | 0.37 ± 0.04 | 0.26 ± 0.04** | 0.26 ± 0.04 | 0.27 ± 0.03 | 0.39 ± 0.05 | 0.26 ± 0.09* |
|  | % of BW | 1.13 ± 0.15 | 0.89 ± 0.17* | 0.77 ± 0.12 | 0.92 ± 0.06 | 1.25 ± 0.19 | 0.96 ± 0.31 |
| **Liver** | Abs (g) | 1.99 ± 0.13 | 1.74 ± 0.28 | 1.7 ± 0.21 | 1.5 ± 0.29 | 1.64 ± 0.13 | 1.4 ± 0.13 |
|  | % of BW | 6.02 ± 0.19 | 5.996 ± 0.87 | 5.04 ± 0.56 | 5.33 ± 1.3 | 5.29 ± 0.28 | 5.22 ± 0.59 |
| **Kidney** | Abs (g) | 0.25 ± 0.03 | 0.23 ± 0.04 | 0.28 ± 0.01 | 0.25 ± 0.02 | 0.23 ± 0.03 | 0.21 ± 0.02 |
|  | % of BW | 0.76 ± 0.07 | 0.80 ± 0.1 | 0.83 ± 0.04 | 0.86 ± 0.02 | 0.74 ± 0.04 | 0.78 ± 0.06 |
| **Pancreas** | Abs (g) | 0.18 ± 0.02 | 0.24 ± 0.04* | 0.22 ± 0.07 | 0.29 ± 0.06 | 0.17 ± 0.04 | 0.23 ± 0.06 |
|  | % of BW | 0.54 ± 0.06 | 0.83 ± 0.15** | 0.64 ± 0.18 | 0.99 ± 0.15<br>(p=0.06) | 0.55 ± 0.1 | 0.86 ± 0.21* |
| **Spleen** | Abs (g) | 0.10 ± 0.02 | 0.09 ± 0.01 | 0.11 ± 0.02 | 0.09 ± 0.02 | 0.1 ± 0.03 | 0.08 ± 0.01 |
|  | % of BW | 0.31 ± 0.08 | 0.29 ± 0.03 | 0.32 ± 0.05 | 0.31 ± 0.04 | 0.31 ± 0.08 | 0.30 ± 0.05 |

Abs: Absolute weight; BW: Body weight; OGT<sup>WT</sup> vs. OGT<sup>CDG</sup> \* $p \leq 0.05$ ; \*\* $p \leq 0.01$ ; OGT<sup>C921Y</sup> line: n = 4 (WT) and n = 6 (C921Y); OGT<sup>N648Y</sup> line: n = 3 (WT) and n = 3 (N648Y); OGT<sup>N567K</sup> line: n = 4 (WT) and n = 4 (N567K)

### Microcephaly and skull deformation in OGT-CDG mice

OGT-CDG patients present with craniofacial dysmorphias, associated with microcephaly in several cases (29). To first investigate whether the OGT^C921Y^ mutation impacts the skull morphology, we performed measurements and microCT imaging on OGT^C921Y^ mice and WT litter mates. Using manual measurements, we observed that OGT^C921Y^ mice exhibited a significant reduction in skull length, while skull width between genotypes was comparable (**Fig. 4A, B**). Analysis of the microCT images revealed that OGT^C921Y^ mice exhibit rounder and smaller skull compared to WT animals (**Fig. 4C**). Superior and lateral views of the skull microCT images were used for two-dimensional measurement of skull parameters using free landmarks (**Fig. 4D**). Overall, 72.2 % of the linear distances were shortened between 2 and 7% in OGT^C921Y^ animals compared to WT litter mates (**Table 2**). Along the rostro-caudal axis, 25.6 % of the distances were significantly reduced (**Table 2**), suggesting mild shortening and deformation of the skull in OGT^C921Y^ mice. The reduction in skull size was associated with a significant decrease in absolute brain weight in OGT^C921Y^ compared to WT mice (**Fig. 4E, F**), suggesting that OGT^C921Y^ mice display a microcephaly phenotype. While we observed no difference in skull length in OGT^N648Y^ and OGT^N567K^ mice compared to their WT littermate (**Fig. S3F, G**), brain weight was reduced (**Fig. S3H, J**). Taken together, these results suggest that OGT-CDG mutations lead to mild skull deformation and microcephaly in mice.

**Figure 4:**
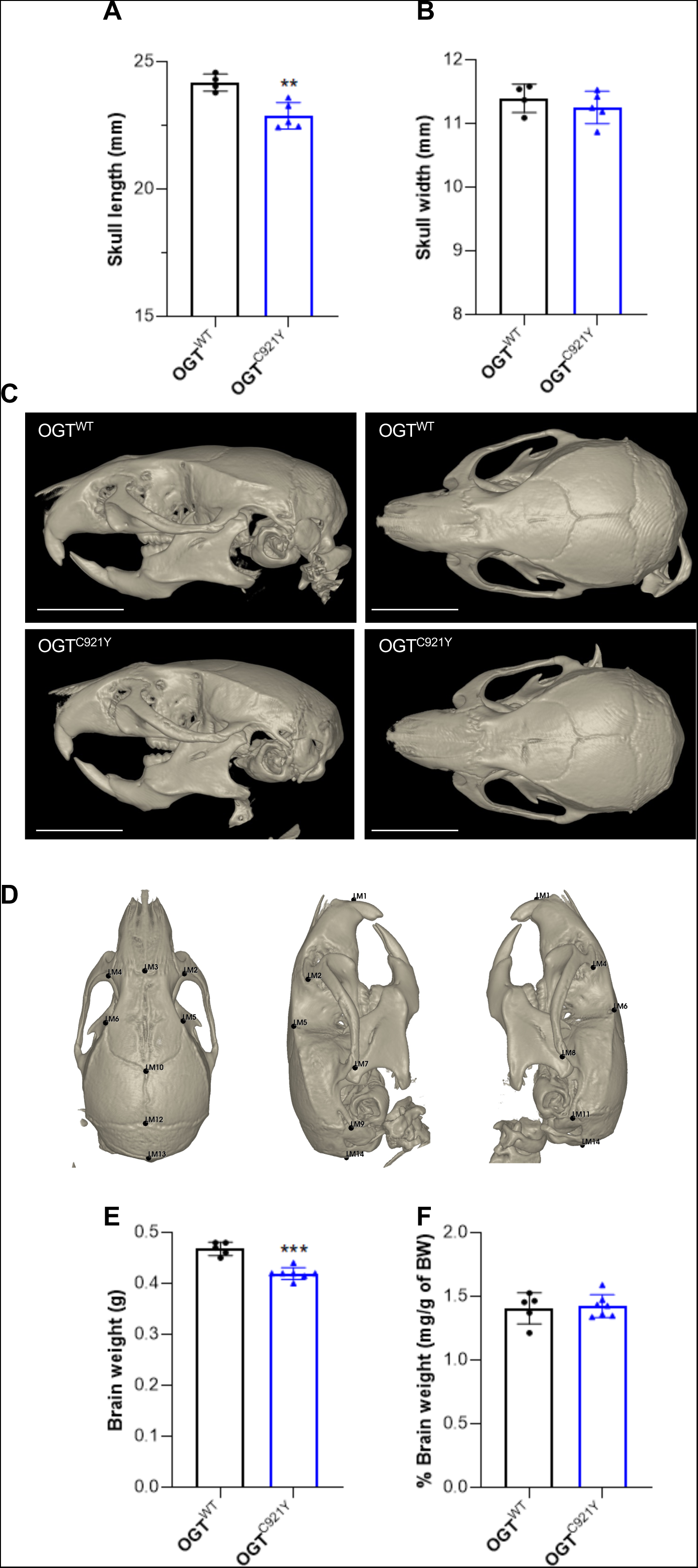
Microcephaly in OGT-CDG mice. Data are represented as mean ± SD. Significance is shown as \**p* < 0.05, \*\**p* < 0.01 and \*\*\**p* < 0.001. **(a)** Measurement of skull length of OGT^WT^ (n = 4) and OGT^C921Y^ (n = 5) mice **(b)** Measurement of skull width of OGT^WT^ (n = 4) and OGT^C921Y^ (n = 5) mice. **(c)** Lateral and superior views of representative 3D reconstruction of the skull of OGT^WT^ and OGT^C921Y^ animals (scale bar = 5 mm). **(d)** Lateral and superior views of a representative 3D reconstruction of a mouse skull presenting 14 landmarks (LM) used for EDMA analysis. **(e)** Absolute brain weight of OGT^WT^ (n = 5) and OGT^C921Y^ (n = 7) mice. **(f)** Percentage of brain weight:body weight ratio of OGT^WT^ (n = 5) and OGT^C921Y^ (n = 7) mice.

**Table 2.** Linear distance ratio of OGT^C921Y^ and OGT^WT^ skulls.

| Landmarks | OGT<sup>C921Y</sup> | p value | Landmarks | OGT<sup>C921Y</sup> | p value | Landmarks | OGT<sup>C921Y</sup> | p value |
| --- | --- | --- | --- | --- | --- | --- | --- | --- |
| 1 to 2 | 0.995 | 0.840 | 2 to 12 | 0.965 | 0.060 | 5 to 14 | 0.964 | 0.014* |
| 1 to 4 | 0.980 | 0.171 | 4 to 12 | 0.977 | 0.229 | 6 to 14 | 0.963 | 0.010** |
| 1 to 3 | 1.000 | 0.953 | 2 to 13 | 0.970 | 0.067 | 7 to 8 | 0.99 | 0.338 |
| 1 to 5 | 0.980 | 0.343 | 4 to 13 | 0.977 | 0.070 | 7 to 9 | 0.942 | 0.009** |
| 1 to 6 | 0.980 | 0.027* | 2 to 14 | 0.962 | 0.004** | 8 to 11 | 0.931 | 0.072 |
| 1 to 7 | 0.976 | 0.165 | 4 to 14 | 0.965 | 0.016* | 7 to 10 | 0.972 | 0.257 |
| 1 to 8 | 0.968 | 0.006** | 3 to 5 | 0.947 | 0.038* | 8 to 10 | 1.009 | 0.255 |
| 1 to 9 | 0.966 | 0.073 | 3 to 6 | 0.957 | 0.224 | 7 to 11 | 0.978 | 0.164 |
| 1 to 11 | 0.957 | 0.015* | 3 to 7 | 0.954 | 0.050* | 8 to 9 | 0.99 | 0.377 |
| 1 to 10 | 0.986 | 0.109 | 3 to 8 | 0.970 | 0.059 | 7 to 12 | 0.945 | 0.057 |
| 1 to 12 | 0.969 | 0.171 | 3 to 9 | 0.962 | 0.010** | 8 to 12 | 0.999 | 0.960 |
| 1 to 13 | 0.970 | 0.076 | 3 to 11 | 0.949 | 0.004** | 7 to 13 | 0.956 | 0.122 |
| 1 to 14 | 0.963 | 0.015* | 3 to 10 | 0.969 | 0.116 | 8 to 13 | 0.991 | 0.411 |
| 2 to 3 | 1.022 | 0.314 | 3 to 12 | 0.951 | 0.014* | 7 to 14 | 0.96 | 0.004** |
| 3 to 4 | 1.045 | 0.172 | 3 to 13 | 0.961 | 0.011* | 8 to 14 | 0.973 | 0.046* |
| 2 to 4 | 1.039 | 0.139 | 3 to 14 | 0.953 | 0.002** | 9 to 10 | 0.996 | 0.829 |
| 2 to 5 | 0.983 | 0.130 | 5 to 6 | 0.984 | 0.492 | 10 to 11 | 0.98 | 0.288 |
| 4 to 6 | 0.991 | 0.647 | 5 to 7 | 0.962 | 0.152 | 9 to 11 | 0.996 | 0.782 |
| 2 to 6 | 1.003 | 0.836 | 6 to 8 | 0.987 | 0.457 | 9 to 12 | 0.985 | 0.771 |
| 4 to 5 | 1.004 | 0.863 | 5 to 8 | 0.992 | 0.400 | 11 to 12 | 1.007 | 0.543 |
| 2 to 7 | 0.950 | 0.040* | 6 to 7 | 0.974 | 0.196 | 9 to 13 | 0.985 | 0.603 |
| 4 to 8 | 0.961 | 0.084 | 5 to 9 | 0.982 | 0.201 | 11 to 13 | 1.014 | 0.326 |
| 2 to 8 | 0.991 | 0.332 | 6 to 11 | 0.955 | 0.046* | 9 to 14 | 0.984 | 0.308 |
| 4 to 7 | 0.984 | 0.389 | 5 to 10 | 0.991 | 0.612 | 11 to 14 | 0.999 | 0.954 |
| 2 to 9 | 0.962 | 0.024* | 6 to 10 | 1.001 | 0.771 | 10 to 12 | 0.936 | 0.278 |
| 4 to 11 | 0.949 | 0.008** | 5 to 11 | 0.971 | 0.034* | 10 to 13 | 0.968 | 0.336 |
| 2 to 10 | 1.000 | 0.771 | 6 to 9 | 0.983 | 0.140 | 10 to 14 | 0.961 | 0.051 |
| 4 to 10 | 1.004 | 0.865 | 5 to 12 | 0.957 | 0.056 | 12 to 14 | 0.967 | 0.351 |
| 2 to 11 | 0.969 | 0.024* | 6 to 12 | 0.971 | 0.246 | 13 to 14 | 0.939 | 0.223 |
| 4 to 9 | 0.981 | 0.199 | 5 to 13 | 0.971 | 0.115 |  |  |  |
|  |  |  | 6 to 13 | 0.976 | 0.170 |  |  |  |

OGT<sup>WT</sup> vs. OGT<sup>C921Y</sup> \* $p \leq 0.05$ ; \*\* $p \leq 0.01$ ; $n = 3$ (OGT<sup>WT</sup>) and $n = 4$ (OGT<sup>C921Y</sup>)

### OGT-CDG mice are hyperactive and show memory defects

The main clinical feature of OGT-CDG is intellectual disability, defined by low IQ (< 70) and maladaptive behaviour (41). We therefore performed a behavioural screen to investigate whether OGT-CDG mice show behavioural or cognitive deficits. We discuss results for the OGT^C921Y^ line, while data for the OGT^N648Y^ and OGT^N567K^ lines are included in the supplemental section (**Fig. S5, S6**). We first investigated general locomotor activity using an open field arena assay. Mice were placed in the arena and allowed to explore for 10 min on three consecutive days to evaluate both exploration of, and habituation to, a new environment. On day 1, there was no difference in distance travelled or speed between OGT^C921Y^ and WT mice suggesting there is no impact of the C921Y mutation on locomotor activity (**Fig. 5A, B**). Interestingly, OGT^C921Y^ mice have higher activity on days 2 and 3, as shown by an increase in distance travelled, speed and time spent mobile compared to WT litter mates (**Fig. 5A-D**).

**Figure 5:**
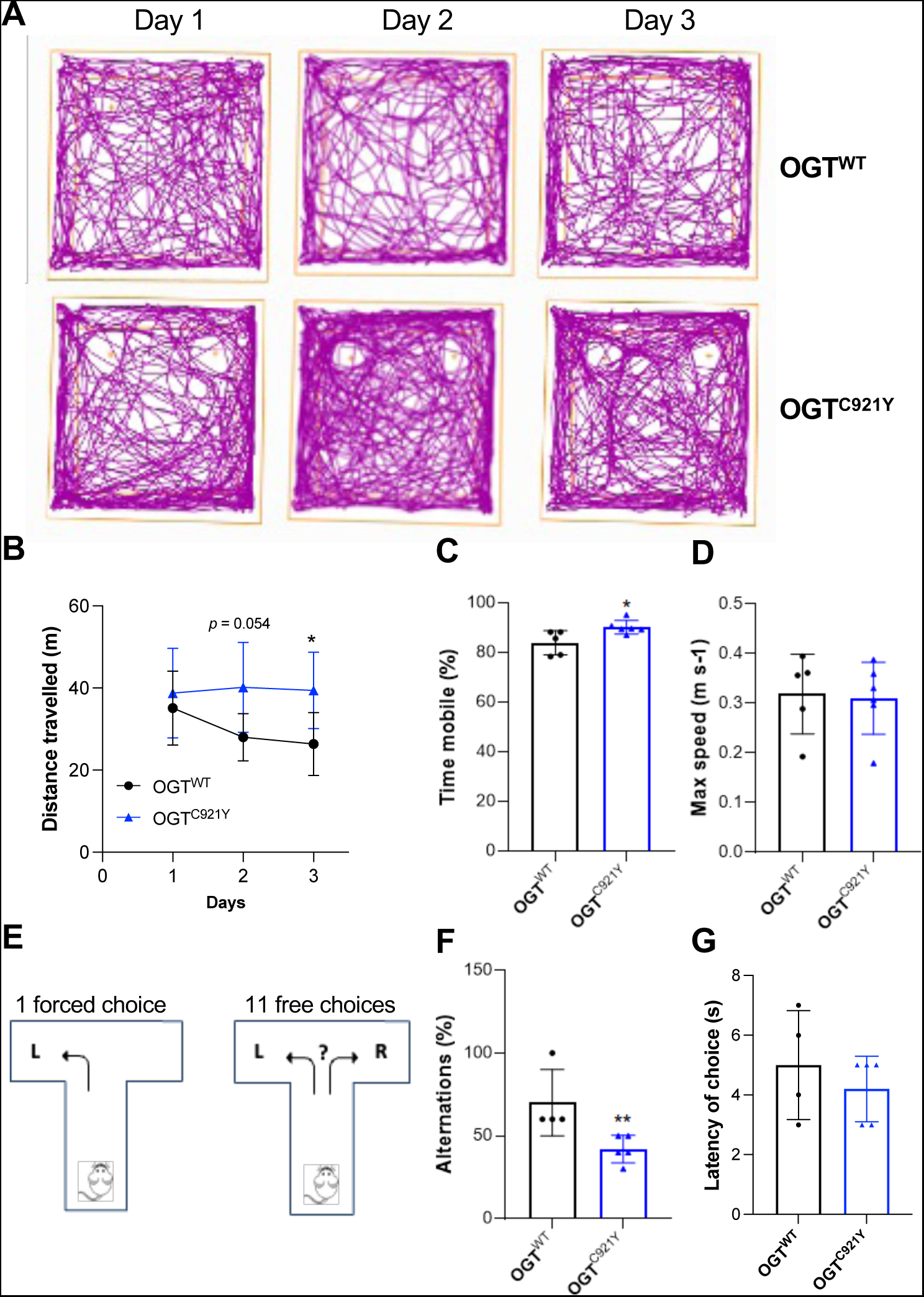
OGT^C921Y^ mice show increased locomotor activity and impaired working spatial memory. Data are represented as mean ± SD. Significance is shown as \**p* < 0.05, \*\**p* < 0.01 and \*\*\**p* < 0.001. **(a)** Representative tracking plot over three consecutive days of an OGT^WT^ (n = 5) and OGT^C921Y^ (n = 6) mice in the open field arena. **(b)** Distance travelled over three consecutive days of OGT^WT^ (n = 5) and OGT^C921Y^ (n = 6) mice in the open field arena. **(c)** Percentage of the mean time spent mobile during three days by OGT^WT^ (n = 5) and OGT^C921Y^ (n = 6) mice in the open field arena. **(d)** Maximal speed displayed by OGT^WT^ (n = 5) and OGT^C921Y^ (n = 6) mice in the open field arena. **(e)** Schematic of the T-maze paradigm. Mice are forced to explore one arm of the maze for the first trial and then are free to explore both arms for eleven additional trials. Each choice and latency were recorded. **(f)** Percentage of correct alternation (L-R-L sequences) of OGT^WT^ (n = 4) and OGT^C921Y^ (n = 5) during the T-maze test. **(g)** Latency to enter in arm of OGT^WT^ (n = 4) and OGT^C921Y^ (n = 5) during the T-maze test.

We next evaluated anxiety, approximated by calculating the fraction of time spent in the periphery versus centre of the arena. Over the three days of testing, OGT^C921Y^ mice demonstrated thigmotaxic behaviour and a preference for the periphery of the arena, as shown by an increase in distance travelled and time spent in the periphery compared to the center of the arena. These findings suggest that OGT^C921Y^ mice may show an anxiety phenotype (**Fig. S4A, B**). To further investigate this, we performed the elevated-plus maze test (EPM) where mice were left to explore a cross shape maze comprised of two closed and two open arms (**Fig. S4C**). Although both WT and OGT^C921Y^ spent more time in the closed arms of the maze, there was no difference in time spent in both areas between OGT^C921Y^ and WT mice (**Fig. S4D**).

We next evaluated motor skills and coordination using grip strength, rotarod and food displacement tests. No differences between WT and OGT^C921Y^ were observed in both the grip strength and rotarod tests, suggesting normal motor and coordination functions in both genotypes (**Fig. S4F, G**). However, in a food displacement test, the OGT^C921Y^ mice removed more food from the apparatus compared to WT mice (**Fig. S4E**), suggesting compulsive behaviour.

Finally, we evaluated spontaneous alternation using a T-maze test as a measure of working spatial memory. Mice were placed in a T-shape maze and allowed to choose between right (R) and left (L) arms for 12 successive trials with the first choice being randomly forced to either left or right. The latency for choosing an arm was similar between OGT^C921Y^ and WT animals **(Fig. 5E).** However, OGT^C921Y^ mice showed a significant reduction in correct L-R-L or R-L-R sequences compared to WT suggesting altered spatial working memory **(Fig. 5F),** a hippocampal-dependent task. Taking together, these results suggest that OGT-CDG mice are hyperactive, display anxiety and show memory defects while possessing normal motor skills.

## Discussion

We have recently described several mutations in *OGT* that are linked to intellectual disability (29). To understand the pathophysiology of this disease, we have generated three independent mouse models each carrying a different catalytically impaired OGT-CDG variant. Although these mutations all cause reduced OGT catalytic activity *in vitro* (30,39,38), OGT^C921Y^, OGT^N648Y^ and OGT^N567K^ animals are viable allowing the phenotypic characterization of OGT-CDG mice and investigation of the function of OGT in neurodevelopment. We have shown that the OGT-CDG mutations lead to a reduction in mass and size as well as altered O-GlcNAc homeostasis in the brain of the mice. In addition, OGT^C921Y^ mice showed microcephaly and cognitive defects reminiscent of clinical observations in OGT-CDG patients (29).

OGT-CDG mice show a reduction in body weight. This was associated with a reduction in body fat mass ratio in OGT^C921Y^ animals compared to WT litter mates. Body weight is determined by a balance between food intake and energy expenditure combining basal metabolism, thermogenesis and physical activity (42). OGT^C921Y^ mice show an increase in locomotor activity that could explain the reduction in body weight observed in these animals. In addition, OGT^C921Y^ show lower levels of blood glucose as well as increase in pancreas size compared to WT. Reduced body weight and pancreas size were also observed in OGT^N648Y^ and OGT^N567K^ lines. Similarly, specific OGT deletion in sensory neurons results in reduced body weight and low blood glucose levels in the *Nav1.8-Ogt^KO^* mouse model (43). Mouse lacking OGT in orexigenic neurons exhibit improved glucose metabolism and are protected from diet-induced obesity (44). Taken together, these studies suggest a role of neuronal OGT in controlling basal metabolism that could affect body composition. Interestingly, *Oga^KO/+^* mice exhibit a lean phenotype similar to our observations in OGT-CDG mice with reduced body fat mass (45). *Oga^KO/+^* animals showed increase in energy expenditure, not caused by elevated locomotor activity but by enhanced thermogenesis through an increase in white to brown adipocytes differentiation (45). Although possible metabolic defects in OGT-CDG mice remain to be further investigated, these observations suggest that in addition to increase locomotor activity, loss of OGT and/or OGA may cause metabolic changes contributing to the lean phenotype observed in OGT-CDG mice.

We observed a reduction in brain size associated with mild skull shortening and deformation in OGT^C921Y^ mice compared to WT animals. The clinical definition of microcephaly is a reduction of head circumference that infers a decrease in brain size (46), we therefore postulate that OGT^C921Y^ mice exhibit a mild microcephaly phenotype. Although microcephaly was not reported in any of the three brothers carrying the OGT^C921Y^ variant (38), it has been observed in patients carrying other OGT-CDG variants including OGT L254F, A259T and R284P variants (47–49). Interestingly, both OGT^N648Y^ and OGT^N567K^ lines show similar reductions in brain size, suggesting that microcephaly may be a more penetrant phenotype in mice than in humans. It is worth noting that no differences in skull length were observed between WT and mutant mice suggesting that the reduction in brain size may have a neurodevelopmental origin rather than skull development defect. The development of the brain depends on several processes including neurogenesis, neuron migration and the balance of proliferation/differentiation of neural stem cells. Although *Ogt* has been shown to be important in all these processes (36,50,51), the mechanisms underlying microcephaly in OGT-CDG remain to be investigated.

We observed behavioural deficits and cognitive defects in OGT^C921Y^ animals. OGT^C921Y^ mice showed increased locomotor activity and exhibit compulsive behaviour in the form of a higher amount of food removed during a food displacement test. Mild hyperactivity has been reported in a patient carrying the OGT^N648Y^ variant (39). Although only two OGT^N648Y^ animals were assessed during the behavioural screening, both mice showed enhanced locomotor activity compared to their WT litter mates, suggesting that hyperactivity may be a penetrant phenotype in OGT-CDG mouse models. Hyperactivity and repetitive/compulsive behaviours are often observed in mouse models recapitulating human ID and autism disorders (52–56). The increase in locomotor activity was not due to enhanced motor capabilities in OGT^C921Y^ animals as no differences were observed in maximum speed in open field, rotarod and grip strength tests. Hyperactivity can be a consequence of animal anxiety in the open field arena. Although OGT^C921Y^ mice showed a preference for spending more time in the periphery over the center of the arena box compared to WT mice suggesting an anxiety-like phenotype, no differences were observed in time/entries in the open arms compared to closed arms during the elevated-maze plus test (EPM). Although both tests are used to evaluate anxiety-like behaviour in mice, they may assess different aspects of anxiety. The open field assay only measures open space-related anxiety whereas EPM includes anxiety induced by open space, light/dark transitions and elevated areas (57). Further experiments will be required to better characterize potential OGT-CDG anxiety phenotypes. Social interaction paradigms, cliff avoidance and novelty tests will be particularly relevant in a context of impaired adaptive behaviour as observed in ID patients.

We have also observed that, unlike the WT mice, the locomotor activity of the OGT^C921Y^ mice does not decrease over the 3 day course of the open field test, suggesting an impairment in spatial habituation learning. Interestingly, defects in habituation learning were also observed in *Drosophila* models carrying OGT-CDG mutations during a light-off jump habituation test (58). As habituation corresponds to the simplest form of learning (59), this suggests that OGT^C921Y^ mice may exhibit learning and memory defects. In addition, OGT^C921Y^ mice show a reduction in spontaneous alternation in a T-maze assay compared to their WT littermates, indicating defects in the spatial working memory process, a hippocampal task. Similarly defects in spatial learning and memory were observed in an inducible neuronal *Ogt^KO^* mouse model (28) suggesting that the loss of OGT activity/protein could mediate the cognitive deficits observed in the OGT^C921Y^ mice.

We did not observe spontaneous seizures events in the OGT^C921Y^ animals that were reported in two of the three brothers affected with the C921Y variant (38). Spontaneous seizures are difficult to study as they usually last a few seconds and may occur infrequently. Proper monitoring will require 24/7 video recording associated with electroencephalograms to detect brain activity in free moving animals (60). Such a set-up has been used to observe myoclonic seizures events in a mouse model of Rett’s syndrome, an ID disorder caused by mutations in *Mecp2* (61). Inducible seizures, either chemically or via sensory stimuli (light/sound), have also been demonstrated in mice carrying *Syngap1* and *Fmr1* mutations associated with ID disorders (62,63). Increased sensibility to light and sounds have been reported in the patient affected with the OGT^N648Y^ variant (39). How OGT-CDG mice may respond to such procedures remains to be explored.

We have previously discussed potential mechanisms underlying the OGT-CDG symptoms (29) and some of these have been assessed in our study. Impaired OGT catalytic activity results in a reduction of global O-GlcNAc levels in the brain of OGT^C921Y^, OGT^N648Y^ and OGT^N567K^ mice. Similarly, hypo-O-GlcNAcylation has previously been observed in cell/*Drosophila* models carrying OGT-CDG variants (30,39). Given that O-GlcNAcylated proteins are particularly abundant in the brain, the loss of O-GlcNAcylation on OGT substrates important for proper brain functioning may contribute to OGT-CDG. Using a filter-based bioinformatics approach combined with structural and clinical data, we have recently predicted the presence of 38 critical O-GlcNAc sites across 22 neuronal proteins already reported to be linked to ID and developmental delay (64). This constitutes a list of potential candidate conveyers for OGT-CDG and further mechanistic dissection of these using OGT-CDG models will help to further understand this disorder. Also consistent with previous findings (30,39), OGA levels were decreased in brain of OGT-CDG mice both at protein and mRNA levels. As OGA has been implicated in intelligence (65) and *Oga* knock-down leads to microcephaly and hypotonia in mice (13), its loss may also contribute to the OGT-CDG phenotype. To investigate this hypothesis, adeno-associated virus vectors (AAVs) or genetic approaches could be used to elevate OGA levels to evaluate whether this could rescue some of the OGT-CDG phenotypes observed in these mouse models. Unexpectedly, OGT protein levels were also reduced in OGT-CDG mouse brains. This has not been previously reported in cell or invertebrate models of OGT-CDG (29,30). O-GlcNAc levels have been demonstrated to regulate *Ogt* transcript levels through alternative splicing (40). However, our data suggest that the decrease of OGT protein levels is not caused by disrupted transcription or alternative splicing. *Ogt* mRNA levels giving rise to the productive form of OGT are in fact increased, a possible compensatory mechanism to maintain global O-GlcNAcylation levels. OGT protein abundance may be disrupted by either protein instability or misfolding leading to proteolytic processing and/or aggregation. Reduction in OGT stability using *in vitro* approaches has been previously reported for OGT-CDG variants (30,49,66,67) but not for OGT^C921Y^ and OGT^N648Y^ mutations (38,39). Some OGT-CDG variants cause conformational changes in the protein that could contribute to OGT misfolding (39,47). Although these studies together suggest that some OGT-CDG mutations could lead to reduced OGT stability, this will require further investigation.

In conclusion we have successfully generated three mouse models of OGT-CDG. Although there are a plethora of lines of investigation to further characterize the phenotypes of these lines, the OGT^C921Y^ mice recapitulate key symptoms observed in OGT-CDG individuals, including cognitive defects and microcephaly. These models will be an invaluable starting point to gain insight into OGT-CDG etiology through identification of underlying mechanisms and conveyer candidates of the disease and provide a platform for evaluation of potential future treatment strategies.

## Materials & Methods

### Generation of OGT-CDG mouse lines and animal husbandry

OGT-CDG mice were produced by microinjection under project licence PPL PB0DC8431 at the Central Transgenic Core of Bioresearch & Veterinary Services at the University of Edinburgh. Female C57BL/6J mice were superovulated with 5IU of Pregnant Mare’s Serum Gonadotropin (PMSG; Prospecbio, hoR-272-b) followed by 5IU of Chorulon (hCG; National Veterinary Services, 804745) 46 h later, and mated overnight with C57BL/6J stud males. Zygotes were harvested at 0.5 dpc. Editing CRISPR reagents were centrifuged through a Millipore filter (UFC30VV25) and injected into the cytoplasm of zygotes on a Zeiss Axiovert 100 using a Femtojet Xpert (Eppendorf), Transferman 4R (Eppendorf) micromanipulators, Vacutip (Eppendorf) holding pipettes and Femtotip (Eppendorf) injection needles. Injected zygotes were cultured to 2-cell stage and then surgically transferred into pseudopregnant Crl:CD1(ICR) or Hsd:ICR (CD-1) females who had been mated with vasectomised Crl:CD1(ICR) the night before. Genomic DNA from offspring was genotyped and sequenced to confirm the presence of the three OGT-CDG mutations. Editing CRISR reagents, Cas9 Nickase (1081062; Integrated DNA Technologies) and primers were purchased from Integrated DNA Technologies and sequences are listed in **Table S1**. Founder OGT-CDG mice were crossed to C57BL/6J WT animals (Charles River UK Limited) for further breeding. Animals were housed in ventilated cages with water and food available *ad libitum* and 12/12 h light/dark cycles. All animal studies and breeding were performed on in accordance with the Animal (Scientific Procedures) Act of 1986 for the care and use of laboratory animals. Procedures were carried under United Kingdom Home Office Regulation (Personal Project Licence PP8833203) with approval by the Welfare and Ethical Use of Animals Committee of University of Dundee.

### EchoMRI and blood sampling

Body composition data from 6-7 months old (OGT^C921Y^) and 6.5-7.5 months old (OGT^N648Y^) male mice was obtained using the EchoMRI^™^ 4in1, in line with the protocol provided by the manufacturer (http://www.echomri.com/Body_Composition_4_in_1.aspx). Basal blood glucose levels from tail vein sampling were measured using Bayer Contour® Glucose Meters and strips.

### MicroCT

55-58 day old mice were euthanised and frozen as intact carcasses and defrosted immediately prior to imaging. Carcasses were imaged at 140 kV and 30 μA using a Nikon XTH 225 ST MicroCT scanner at 50 micron resolution. Two-dimensional images were used to generate 3D volumes using 3D slicer (https://www.slicer.org/) (68). Coordinates of 14 landmarks on the skull were recorded blindly from three-dimensional CT images for analysis of the skull morphology.

### Tissue collection and disruption

Brain tissues from 80-91 (OGT^C921Y^), 86-90 (OGT^N648Y^) and 132-134 (OGT^N657K^) days old male mice were rapidly dissected, snap frozen in liquid nitrogen, and stored at −80 °C. Tissues were disrupted in Phosphate-Buffered Saline two times at 5000 rpm for 30 s with 10 s break using Precellys^®^ 24 Touch homogenizer (Bertin Technologies). Homogenates were split in half for further protein and RNA extractions.

### Western blot

Brain homogenates were lysed using 10x RIPA buffer (Cell Signaling), centrifuged at 14,000 rpm for 20 min at 4 °C, and the protein concentration was determined with Pierce 660 nm protein assay (Thermo Scientific). Proteins were separated on precast 4-12% NuPAGE Bis–Tris Acrylamide gels (Invitrogen) and transferred to nitrocellulose membrane. Membranes were incubated with primary antibodies in 5% bovine serum albumin in Tris-buffered saline buffer with 0.1% Tween-20 overnight at 4 °C. Anti-OGA (1:500 dilution; HPA036141; Sigma), anti-O-GlcNAc (RL2) (1:500 dilution; NB300-524, Novus Biologicals), anti-OGT (F-12) (1:1000 dilution; sc-74546; Santa Cruz), mouse anti-lamin B (1:10000 dilution; 66095-Ig; Proteintech) and rabbit anti-lamin B (1:5000; 12987-1-AP; Proteintech) antibodies were used. Next, the membranes were incubated with IR680/800-labeled secondary antibodies (Li-cor) at room temperature for 1 h. Blots were imaged using a Li-Cor Odyssey infrared imaging system (Li-Cor), and signals were quantified using Fiji software. Results were normalized to the mean of each corresponding WT replicates set and represented as a fold change relative to WT.

### qPCR analysis

Total RNA was purified from brain homogenates using RNAeasy Kit (Qiagen), and then 0.5 to 1 µg of sample RNA was used for reverse transcription with the qScript cDNA Synthesis Kit (Quantabio). Quantitative PCR reactions were performed using the PerfeCTa SYBR Green FastMix for iQ (Quantabio) reagent, in the CFX Connect Real-Time PCR Detection System (BioRad), employing a thermocycle of one cycle at 95 °C for 30 s and then 40 cycles at 95 °C for 5 s, 60 °C for 15 s, and 68 °C for 10 s. Data analysis was performed using CFX Manager software (BioRad). Samples were assayed in biological replicates with technical triplicates using the comparative Ct method. The threshold-crossing value was normalized to internal control transcripts (Gapdh, Actb, and Pgk1). Primers used for qPCR analysis are listed in Table S2. Results were normalized to the mean of each corresponding WT replicate set and represented as a fold change relative to WT.

### Behaviour

For behavioral testing, male animals were transferred in standard cages before the start of the study. Prior to any behavioural testing, all animals experienced daily handling by the experimenter for one week. All handling took place in the experimental room at the same time each day. All animals were housed in groups of 2 to 6 individuals. Animals were 10–13 weeks old at the start of behavioural testing, reaching 16–19 weeks old at the end. The experiments were performed blind, with the experimenters unaware of genotype while performing the test and analysing the data. *Open Field:* The activity of mice in an open-field maze was recorded using Any-maze video tracking software. Individual animals were placed in a 40 x 40 x 40 cm opaque box and were allowed to explore for 10 min over three consecutive days. Time spent in the centre and periphery and time moving in the periphery were analysed to investigate parameters of locomotion and anxiety. *Spontaneous alternation:* The spontaneous alternation test was performed in a T-maze and was used to assess spatial working memory. For the first trial, individual mice were placed in the starting area and forced to explore one arm randomly assigned for 1 min. Then mice were free to choose left or right arms for another 11 trials. Each arm entry was recorded to calculate the percentage of spontaneous alternation corresponding to the number of correct Left-Right-Left or Right-Left-Right sequences. Mice that were able to remember which arms they had entered most recently would choose a different one to explore. *Elevated-plus maze:* Anxiety-like behavior was assessed using elevated plus maze paradigm. Mice were placed in the center of the cross-shape maze comprised of two open arms and two closed arms and allowed to explore the maze for 15 min. Time, distance and number of entries in each open and closed arms were analyzed to investigate the level of anxiety. *Rotarod and grip strength:* Coordination and gross motor skills were assessed using rotarod and grip strength tests. Mice were placed on a Ugo Basile 7650 rod with an increase in speed from 4 to 40 rpm every 30 sec for 5 min. Grip strength was assessed using a familiar inverted cage grid to measure hanging capability. Latency to fall from both rotarod and lid were measured.

### Statistics

Statistical analyses were performed with Prism 9. D’Agostino & Pearson, Shapiro–Wilk, and Kolmogorov-Smirnov normality tests were performed to verify normality. For data that fulfilled normality requirements, unpaired *t* test was used for pairwise comparisons of wild-type and OGT-CDG data. For data sets that did not fulfill normality, Mann-Whitney test was used for pairwise comparisons of wild-type and OGT-CDG data.

## Acknowledgements

This work was funded by a Wellcome Trust Investigator Award (110061) and a Novo Nordisk Foundation Laureate award (NNF21OC0065969) to D.M.F.v.A. We thank Dr. Stewart J. Chalmers from Aberdeen University for assistance with MicroCT data collection. We also thank Conor Mitchell for constructive feedback.

## Author contributions

F.A., A.M., and D.M.F.v.A conceived the study; F.A., N.O., A.M and D.M.F.v.A. performed experiments; A.T.F. performed molecular biology; F.A., N.O., A.M. and D.M.F.v.A. analysed data and F.A., A.M. and D.M.F.v.A. interpreted the data and wrote the manuscript with input from all authors.

## Conflict of interest

No conflict of interest.

## Notes

### Competing Interest Statement

The authors have declared no competing interest.

