## Supplementary figures and images for "Neurodevelopmental defects in a mouse model of O-GlcNAc transferase intellectual disability"

### Supplemental material

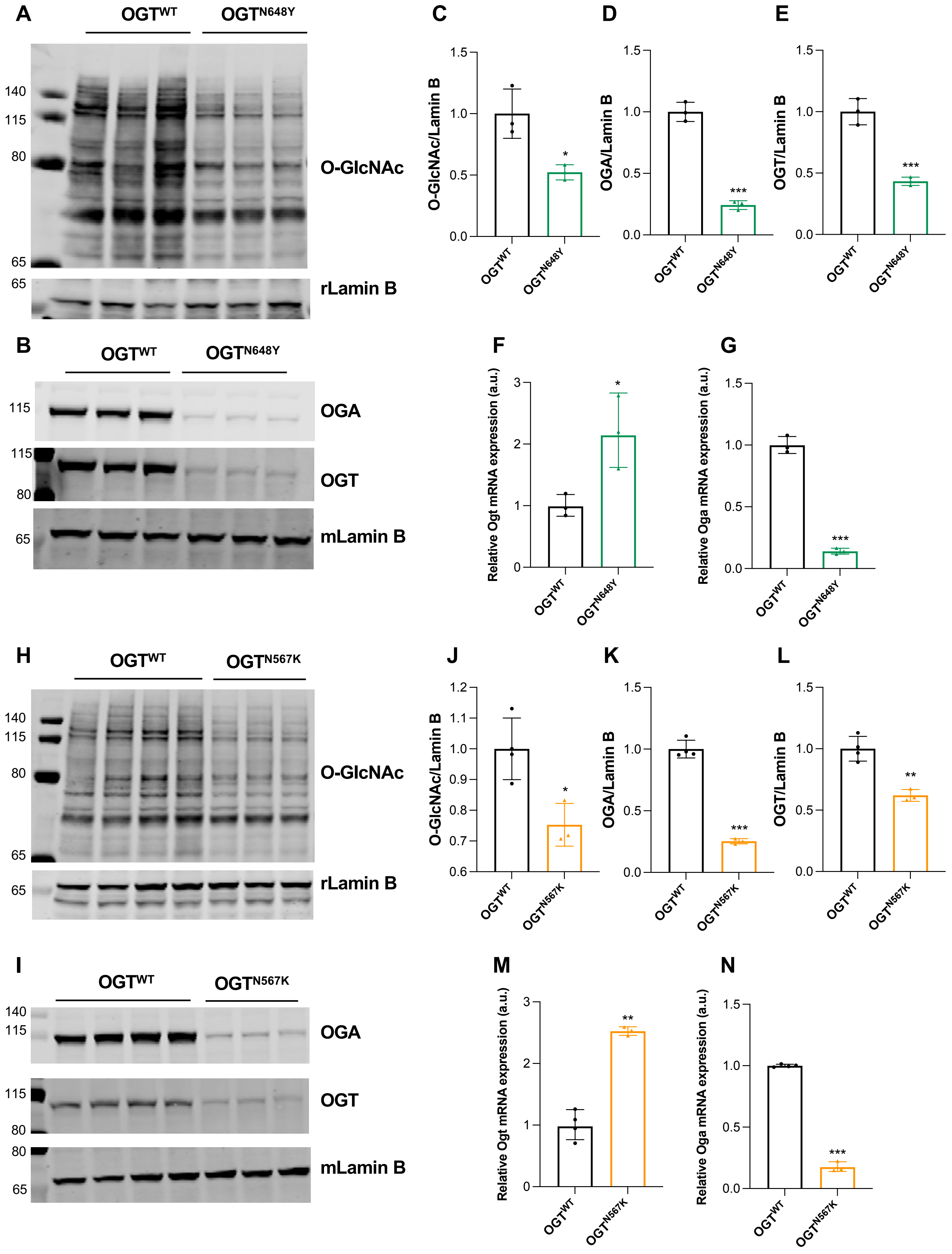

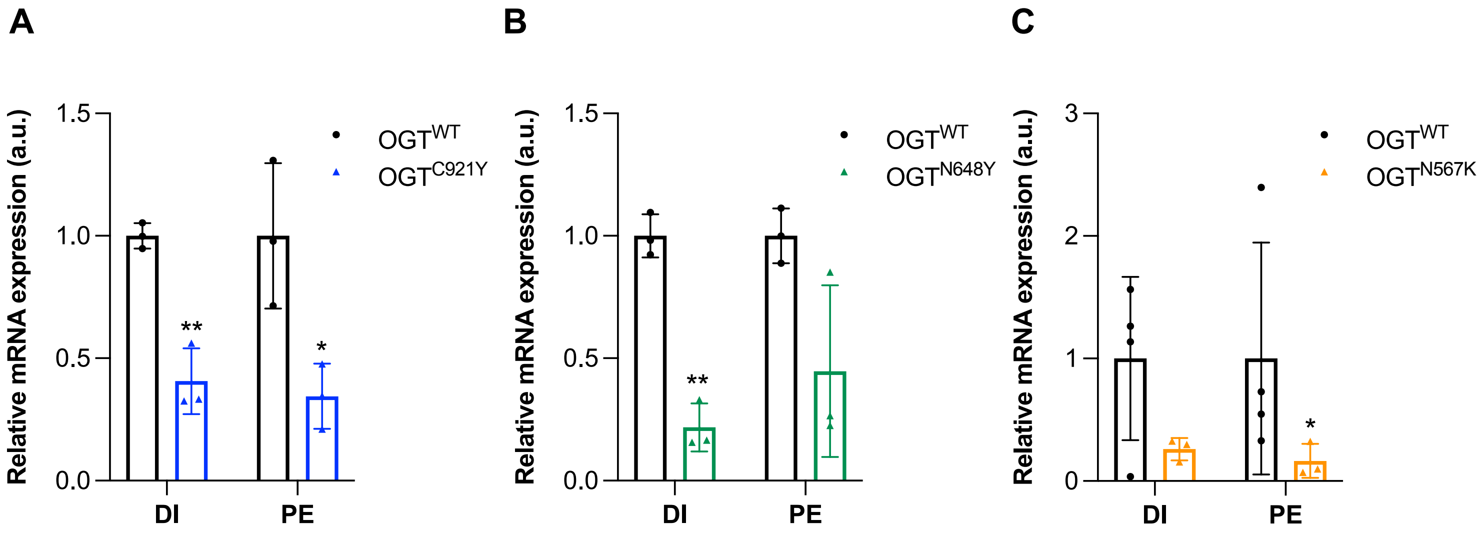

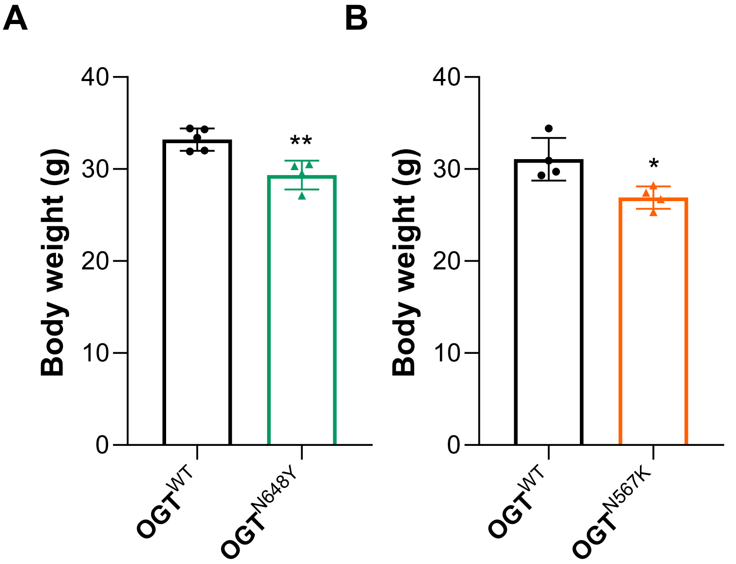


C

| | OGT^WT | OGT^N648Y |
|---|---|---|
| Fat mass (g) | 13 | 3.1 ± 1.3 |
| Fat mass:BW ratio (%) | 28 | 9.3 ± 2.7 |
| Lean mass (g) | 32 | 27 ± 3 |
| Lean mass:BW ratio (%) | 67 | 84 ± 1.4 |
| Glycemia (mmol.L$^{-1}$) | 6.7 | 6.3 ± 2.5 |

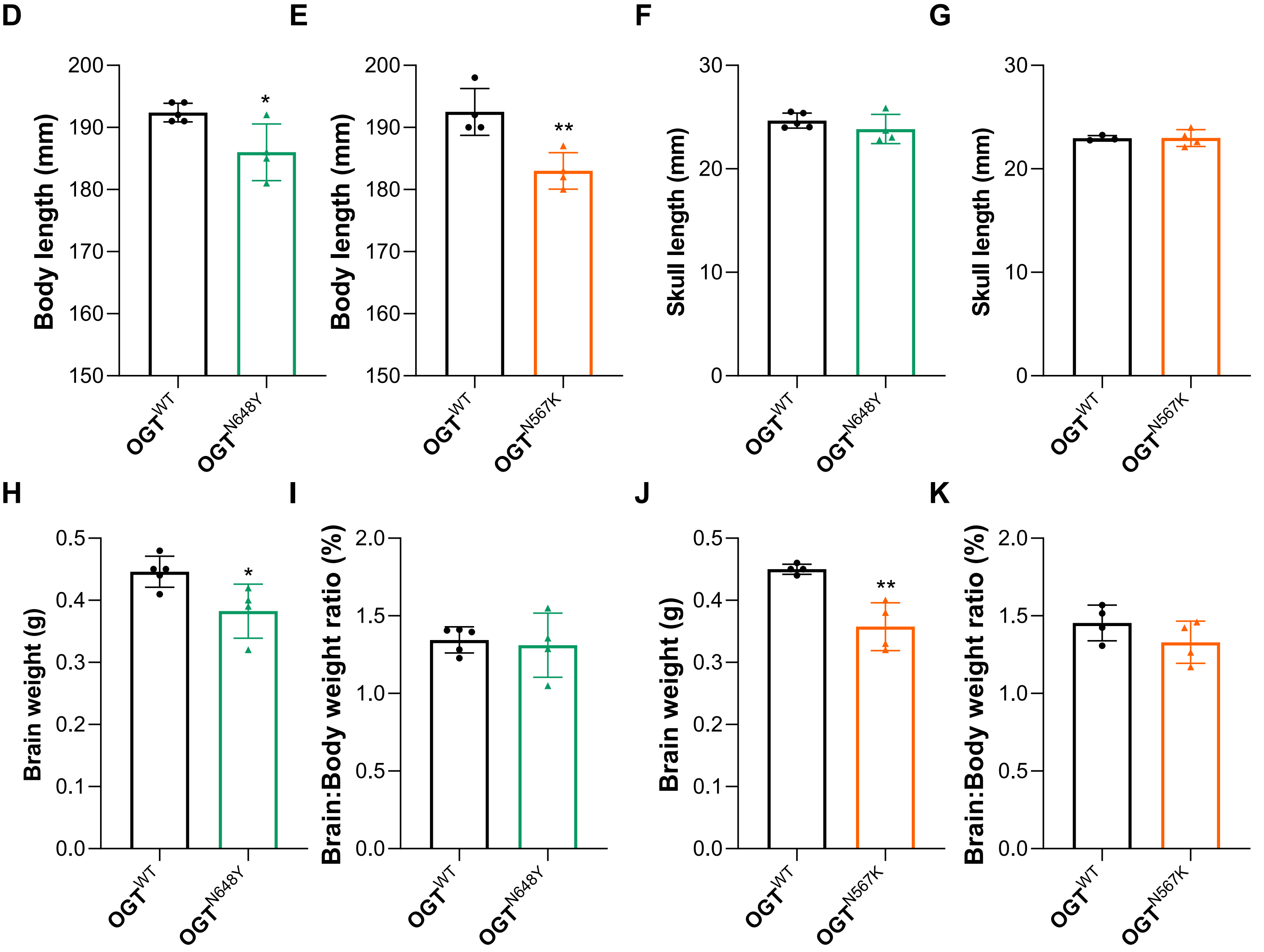


S3

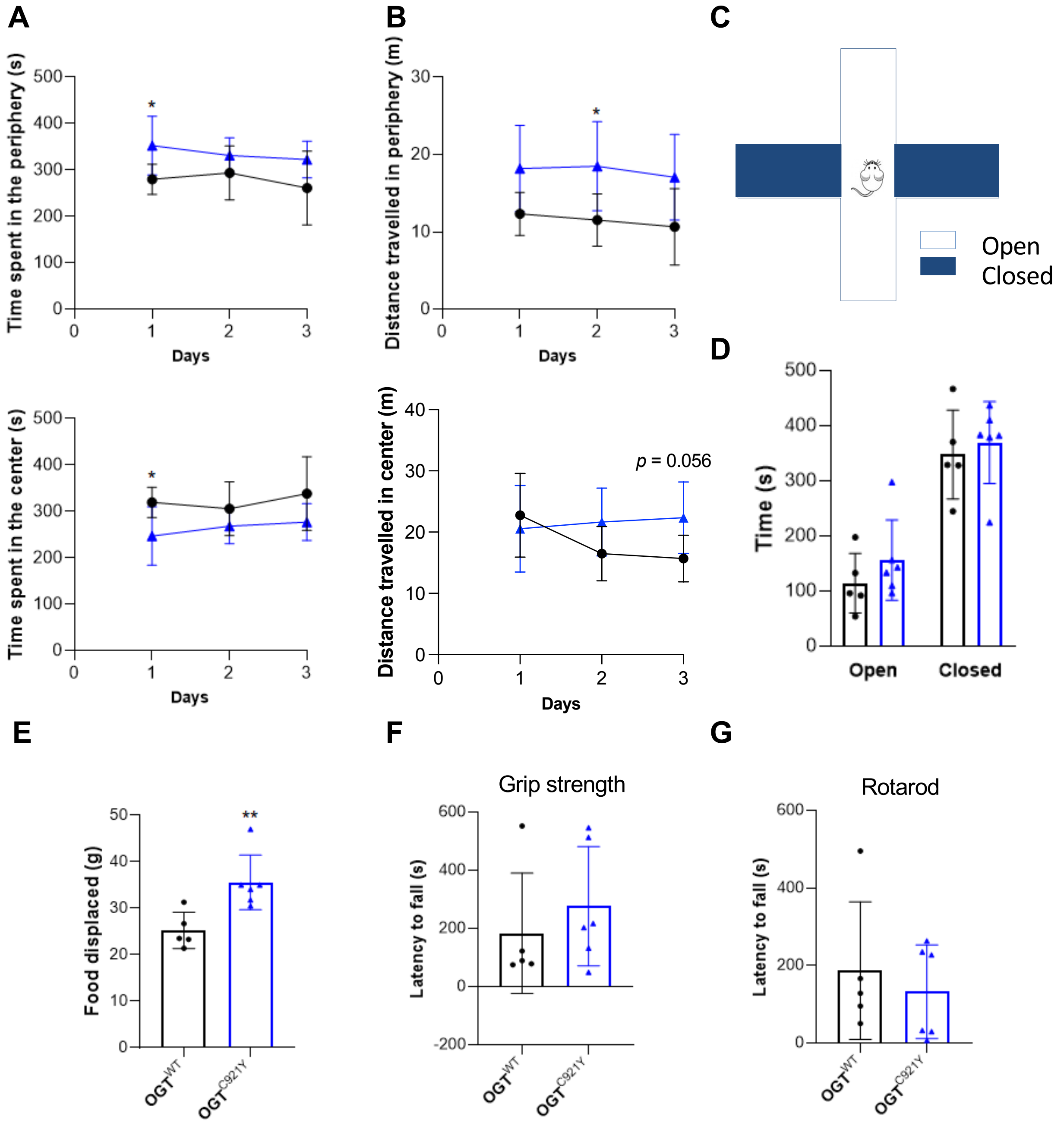

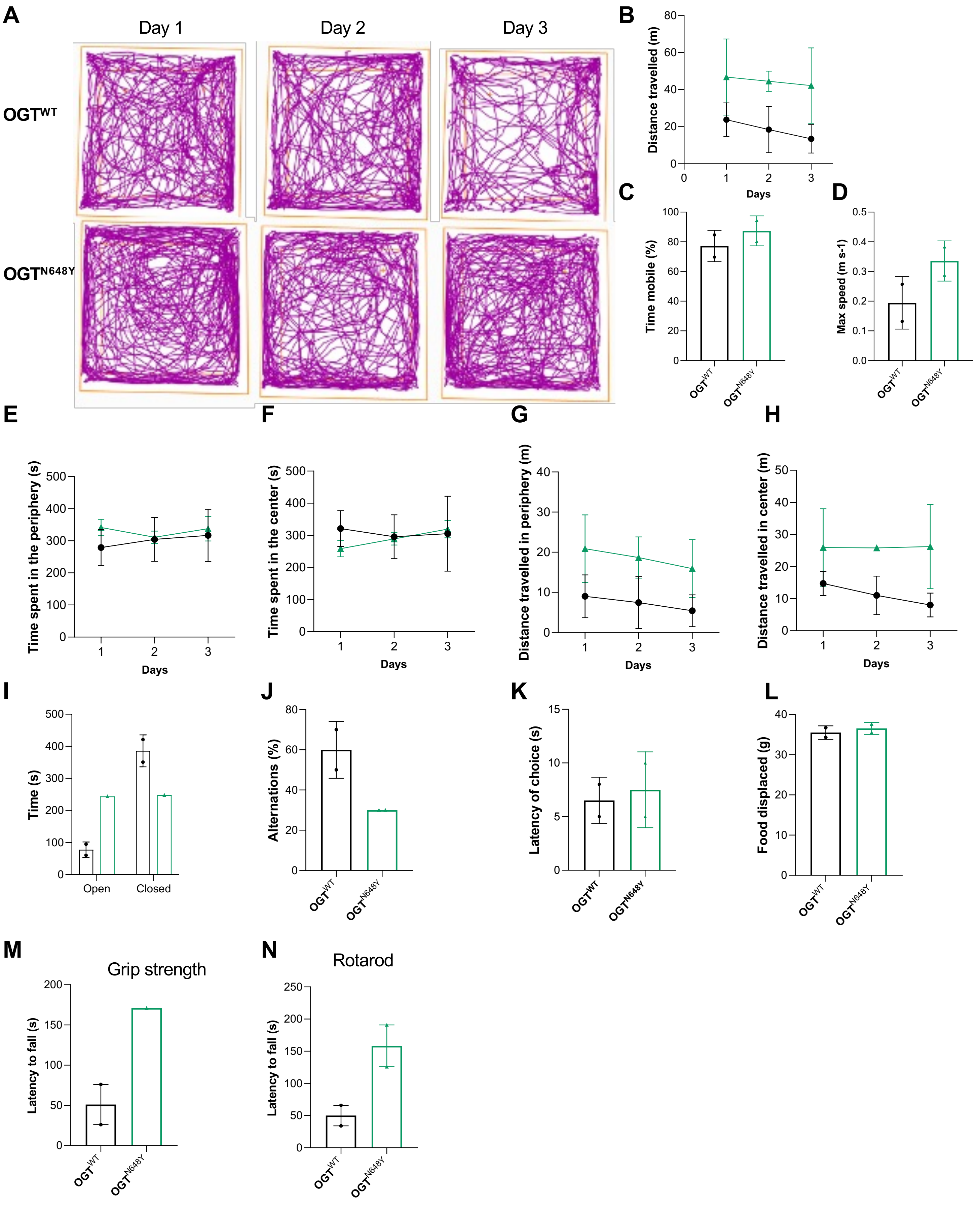

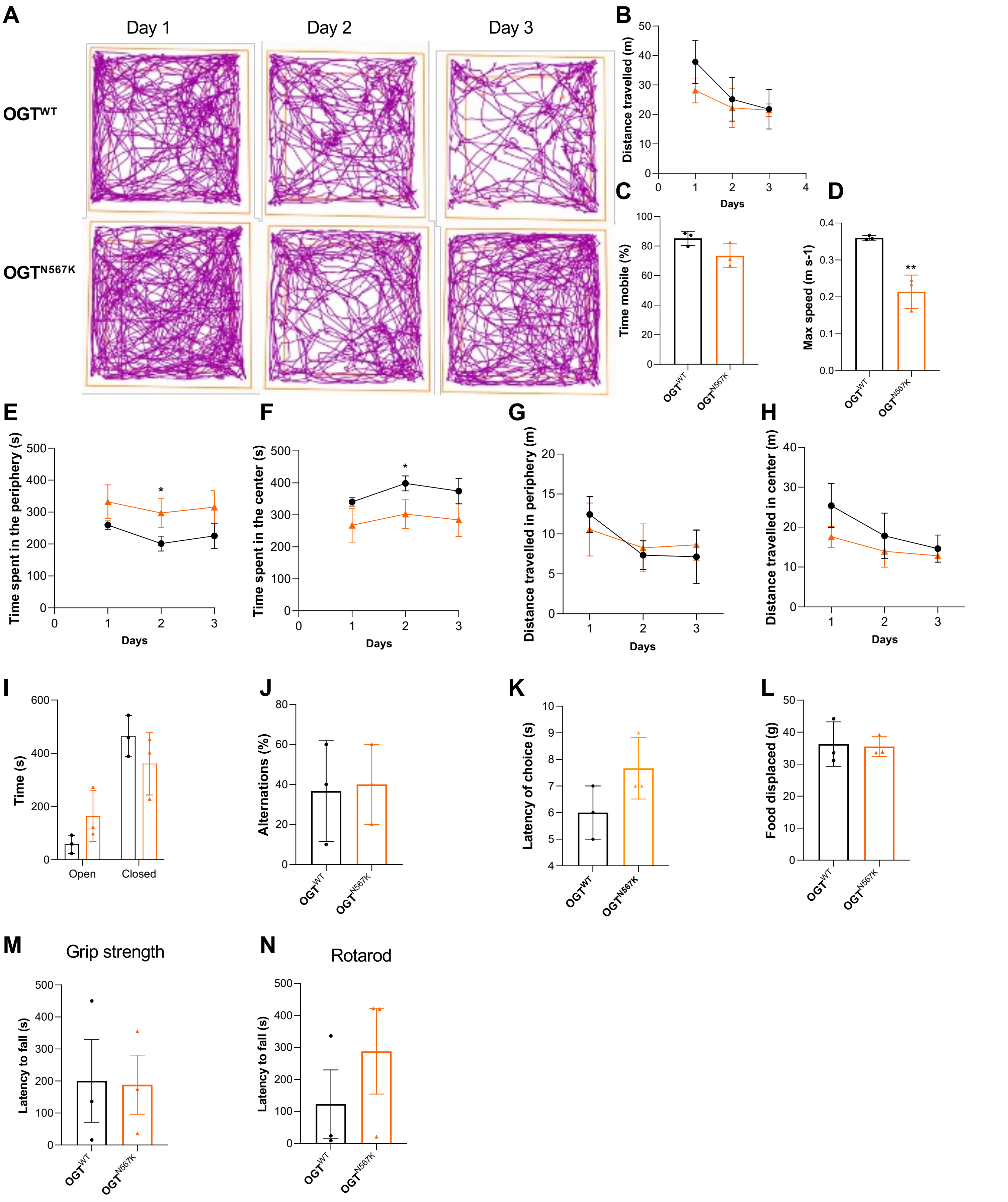
